# A Novel Method for Quantifying Traction Forces from Hexagonal Micropatterned Features on Deformable Poly-Dimethyl Siloxane Sheets

**DOI:** 10.1101/479790

**Authors:** Brian P. Griffin, Christopher J. Largaespada, Nicole A. Rinaldi, Christopher A. Lemmon

## Abstract

Many methods exist for quantifying cellular traction forces, including traction force microscopy and microfabricated post arrays. However, these methodologies have limitations, including a requirement to remove cells to determine undeflected particle locations and the inability to quantify forces of cells with low cytoskeletal stiffness, respectively. Here we present a novel method of traction force quantification that eliminates both of these limitations. Through the use of a hexagonal pattern of microcontact-printed protein spots, a novel computational algorithm, and thin surfaces of polydimethyl siloxane (PDMS) blends, we demonstrate a system that quantifies cellular forces on a homogeneous surface that is stable, easily manufactured, and can quantify forces without need for cellular removal.

## Introduction

Cellular contractile forces are generated by myosin motors pulling on filaments of the cytoskeletal protein actin; these forces are transmitted outside the cell via a combination of both adherens junctions (cell-cell contact) and focal adhesions (cell-extracellular matrix (ECM) contact). Cell-ECM forces, or cellular traction forces, affect not only the contractile and migratory state of the cell, but also affect cellular programs such as differentiation and proliferation (1–9). Several methods exist to quantify cellular traction forces generated by cells. However, each of these has limitations.

One such method is traction force microscopy, in which deformable sheets of silicone rubber are seeded with fluorescent particles that move under cell deflection (10). This method requires the application of boundary conditions and constraints to solve for cell-generated forces, which can bias the solution greatly. Additionally, this method requires that cells be detached at the conclusion of an experiment to determined the unstressed state of the silicone rubber sheet. Variations of this protocol have been developed in which substrates are patterned with microcontact-printed proteins in specific locations, which eliminates the need to make assumptions about the location of traction forces (11). However, these assays still require that cells be removed in order to determine undeflected positions of the substrate.

Cellular traction forces can also be quantified through the use of microfabricated post arrays (12), which consist of an array of deformable micropillars that bend under cell-applied force. The discrete nature of the posts can cause cells with low cytoskeletal stiffness to sag between adjacent posts, which leads to inaccurate quantification of forces. Additionally, the discrete nature of the pillars creates a scenario in which strains are not propagated throughout the surrounding substrate, which has been identified as a key aspect of cell migration and durotaxis (13).

Here we present a novel analysis and experimental approach to traction force quantification. By coupling low elastic modulus blends of polydimethyl siloxane (PDMS) with a hexagonal microcontact printing pattern and a novel finite-element analysis approach, we have developed a system that overcomes previous limitations and is easily adoptable by labs with no history or experience in traction force microscopy. The new system: a) does not require removal of cells to calculate the undeflected state, b) is a continuous substrate, which allows strains to propagate throughout the system; and provides improved data for traction forces generated by cells with reduced cytoskeletal integrity.

## Methods

### Protocol Summary

Steps for the protocol are described in detail below. The summarized protocol is described here and is shown visually in Fig. 1:

**Fig. 1.**
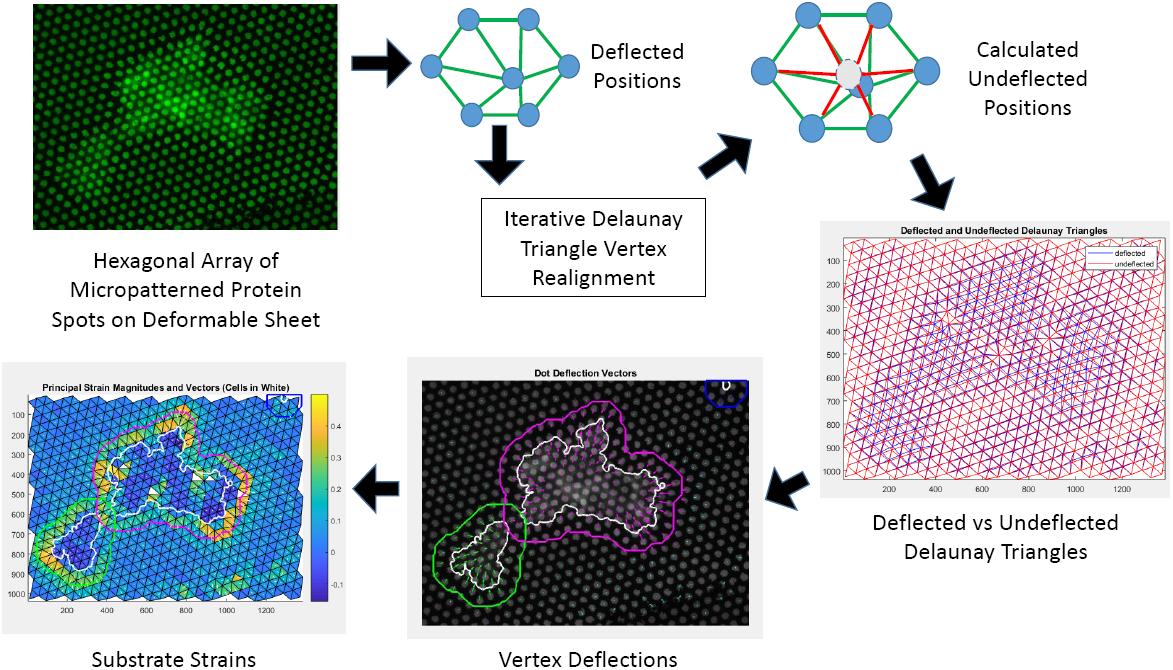
Methodology and Approach for Traction Force Measurement. Deformable polydimethyl siloxane (PDMS) substrates are microcontact printed with a hexagonal array of protein spots. Cells deflect the spots, and a Delaunay triangulation is generated from the deflected spots. Delaunay triangle vertices are iteratively shifted to the center of the surrounding six spots in order to calculate the undeflected positions. Deflections are used to calculate principal strains and principal strain directions for each Delaunay triangle

#### 1) PDMS substrate fabrication

Glass coverslips were spin coated with a blend of PDMS that is roughly 3 orders of magnitude softer than Sylgard 184 PDMS, which facilitates deflections in the substrate under cell-applied traction forces.

#### 2) Substrate patterning

Using a previously published “stamp-off” technique (14), we created a PDMS stamp that contained the desired hexagonal pattern of circular protein spots. This stamp was used to microcontact-print the desired hexagonal pattern onto PDMS substrates Fig. 2A. A representative patterned surface is shown in Fig. 2B. Pluronics F127 was used to limit cell adhesion to the printed pattern.

**Fig. 2.**
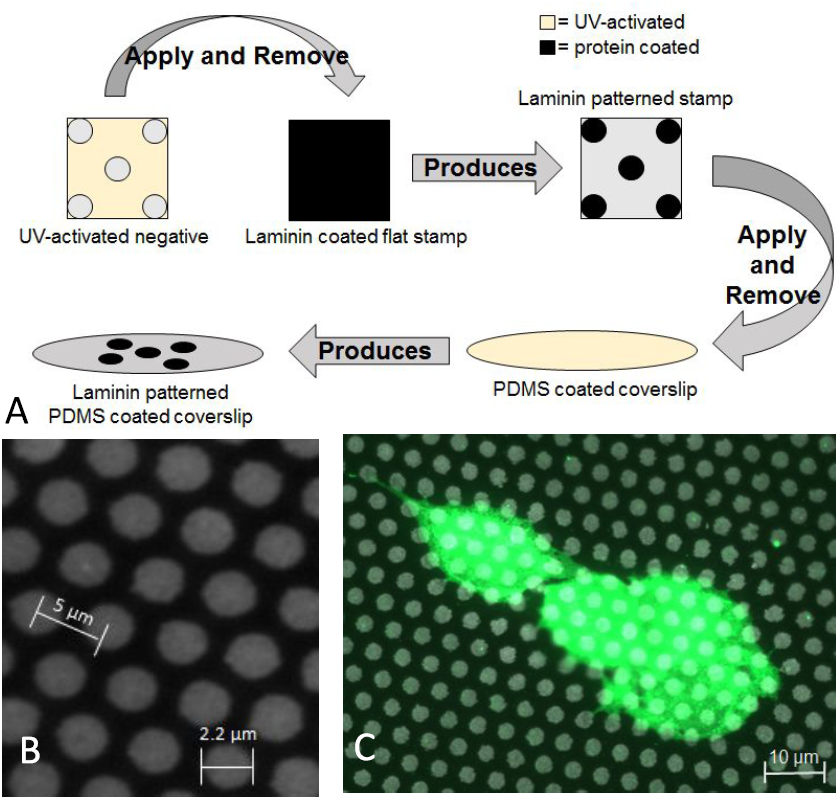
Hex Force pattern generation. A) Schematic of patterning protocol. A surface patterned with features that are the inverse of the desired pattern is treated with UV ozone and applied to a laminin-coated PDMS stamp. This results in a PDMS stamp with the desired pattern. This stamp is then applied to a UV ozone-activated PDMS-coated coverslip. B) A representative image of the laminin-patterned surface. C) A representative image of cells (green) attached to the laminin-patterned surface.

#### 3) Cell plating and imaging

Cells were plated onto the patterned surfaces, and images of the deflected patterns were acquired via immunofluorescence microscopy (Fig. 2B,C).

#### 4) Analysis: Delaunay Triangulation

Centroids of each spot were identified using image processing tools in MATLAB (Fig. 3A). The hexagonal pattern of microcontact printed circles was used to generate a Delaunay triangulation (Fig. 3C). This pattern was utilized to both calculate the undeflected positions of circles as well as to discretize the substrate for strain analysis.

**Fig. 3.**
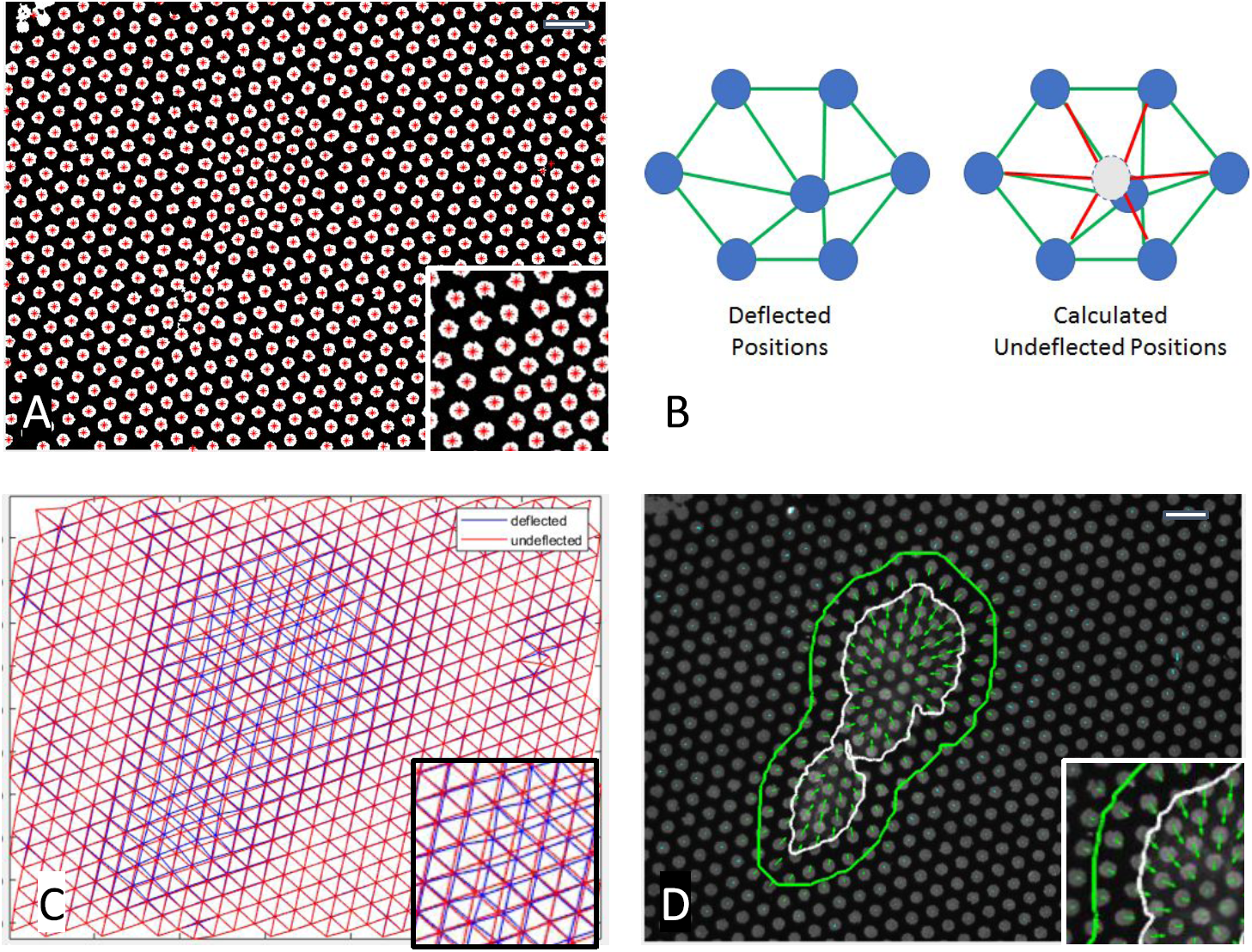
Algorithm to calculate cell-driven deflections of patterned surfaces. A) Protein spots are microcontact printed onto a PDMS surface in a hexagonal pattern; cells attached to the surface cause microcontact printed spots to deflect. Spot centroid is identified via an image processing algorithm in Matlab (+). B) For each identified protein spot, the HexForce analysis package iteratively moves the protein location to the center of its 6 neighbors, until a converged solution is reached. This indicates the undeflected position of each spot. C) Delaunay triangulation of the measured deflected positions (blue) and the calculated undeflected positions (red). D) Deflected and undeflected nodes in C are used to calculate displacement vectors (green arrows). Cell outline is shown in white, and the region of influence for which the cell generates deflections is shown in green. Scale bar equals 10 *µ*m.

#### 5) Analysis: Identification of Missing and Mis-identified Spots

Voronoi diagrams were generated to identify degenerate hexagonal patterns in the images (Fig. 4A), which were used to replace missing or misidentified circles in the images.

**Fig. 4.**
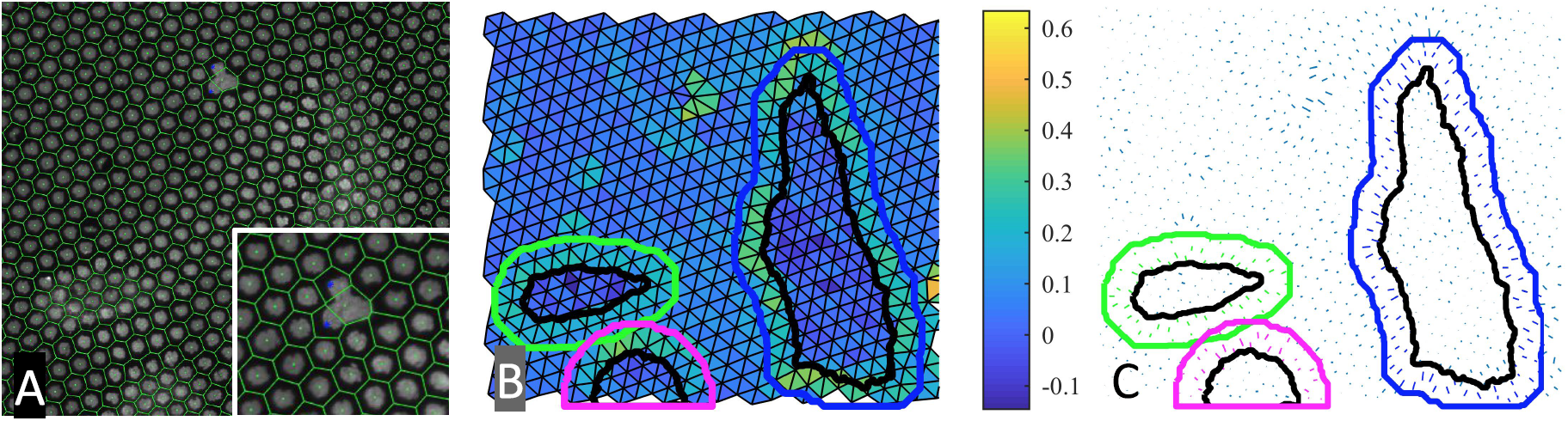
Correction for mis-identified spots and calculation of substrate strain. A) A Voronoi diagram is generated for hexagonal patterns (green); mis-identified spots are determined by examining degenerate geometries in the pattern, and replacement spots are added (blue). B) Principal strains for each Delaunay triangle are calculated using the deflection map and linear plane strain equations. Cell outlines are indicated in black, while the region of influence surrounding each cell is colored per cell. Triangles are colored with principal strain magnitude. C) Principal strain vectors for Delaunay triangles. Strains are shown as vectors whose length corresponds to the magnitude of strain and whose direction corresponds with the direction of principal strain.

#### 6) Analysis: Calculation of Undeflected positions

An iterative image processing algorithm was performed, in which each dot centroid was moved to the average of its 6 neighbors (Fig. 3B). Iterations proceeded until the array of dot centroids no longer changed, indicating that each dot was at the center of its neighbors, as would be expected in the undeflected pattern (Fig. 3C). Deflections were calculated by quantifying the vector from the original position to the calculated undeflected position (Fig. 3D).

#### 7) Analysis: Principal strains and stresses from dot deflections

Deflections at the Delaunay triangle corners were used to calculate plane strain in each triangle (Fig. 4B, C). These strains were transformed to calculate the magnitude and direction of principal strain and stress on each triangle. Plane stress was integrated over all triangles to determine total cell-generated force.

### PDMS Substrate Fabrication and Preparation

PDMS is frequently used in biological applications due to its biocompatibility and straightforward polymerization. Typical biological applications use a 10:1 base:crosslinker ratio of Sylgard 184 PDMS (Dow Chemical), which has an elastic modulus of roughly 2 MPa, which is too rigid for sheets to deflect in response to cell-generated traction forces. Here we utilized a blend of 20:1 Sylgard 527:Sylgard 184. Sylgard 527 has an elastic modulus of roughly 5 kPa, which is more physiologically relevant and will deflect under cell traction forces. The two formulations were blended to give a polymer with an elastic modulus of approximately 7.5 kPa, which created a surface that deflected under cell traction forces, but was also sufficiently rigid to allow for microcontact printing and subsequent processing. Previous studies have demonstrated that 527:184 blends are biologically compatible and create a homogeneous substrate (15). We first generated a 1:1 base:crosslinker ratio of Sylgard 527 and a 10:1 base:crosslinker ratio of Sylgard 184. Each individual polymer was degassed under vacuum, and then combined at the 20:1 ratio. All ratios are mass:mass.

PDMS blends were then spin coated onto 25 mm diameter glass coverslips at 3000 RPM for 35 seconds using a Laurell Technologies WS-650MZ-23 spin coater to generate a roughly 10 *µ*m thick uniform layer. Coated coverslips were cured overnight at 110 °C.

Photolithographic techniques were used to generate an array of cylindrical micron-scale pillars on the surface of a silicon wafer as previously described (16). Briefly, silicon wafers were spin-coated with a layer of UV-crosslinkable SU-8 negative photoresist. The wafer was then brought into conformal contact with a quartz mask that was patterned with a hexagonal array of 2 *µ*m circles with a 5 *µ*m center-to-center spacing. The wafer was exposed to UV light through the mask in order to crosslink the SU-8. Un-crosslinked SU-8 was removed via standard photolithography techniques to create an array of 2 *µ*m diameter by 5 *µ*m tall pillars on the surface.

Wafers containing the micropillar arrays were coated with 10:1 base:crosslinker Sylgard 184, cured overnight and removed to create inverse patterns necessary for the “Stamp-Off” technique originally described by Desai, et. al. (14). These inverse patterns were exposed to UV ozone for 7 minutes and brought into conformal contact with a square block of 10:1 base:crosslinker Sylgard 184 that had been incubated with 1 *µ*g/mL laminin for 10 minutes at room temperature. UV ozone exposure created reactive oxygen groups on the surface of the inverse pattern, and conformal contact with the laminin-coated surface removed laminin from these regions (Fig. 2A).

The laminin-coated stamp, which contained the appropriate hexagonal pattern of laminin, was brought into conformal contact with PDMS-coated glass coverslips that had been exposed to UV ozone for 7 minutes. This transferred the hexagonal pattern to the PDMS-coated coverslip. To ensure that cell adhesion was limited to the microcontact printed pattern, coverslips were incubated with 1% Pluronics F127 in phosphate-buffered saline (PBS) for 30 minutes at room temperature. A representative image of the printed hexagonal pattern of laminin circles is shown in Fig. 2B.

### Immunofluorescence Labeling and Microscopy

Cells were seeded onto laminin micropatterned PDMS coverslips and incubated at 37°C for 24 hours. Cells were then rinsed in 1X PBS and fixed using 4% paraformaldehyde in PBS for 20 minutes. Cells were immunofluorescently labeled to detect patterned laminin, actin, and nuclei using the antibodies and reagents listed below. Prior to both primary and secondary antibody incubation, permeabilized cells were incubated with 0.1% Bovine Serum Albumin (BSA) for 5 minutes to prevent non-specific antibody reactivity. Cells were incubated at 37°C for 30 minutes with primary antibodies, washed in 0.1% BSA, and subsequently incubated at 37°C for 30 minutes with secondary antibodies. Immunofluorescence images were taken using a Zeiss AxioObserver Z1 inverted fluorescence microscope with ZEN 2011 software and processed through an original MATLAB code described below. Images were acquired using a 63x objective.

### Cells and Reagents

Both 3T3 mouse embryo fibroblast cells and MCF10A human mammary epithelial cells were used to validate the system. 3T3 cells (ATCC) were cultured at 37°C with 5% CO2 in Dulbecco’s Modified Eagle Medium (DMEM) containing 4.5 mg/mL glucose (Gibco), 10% fetal bovine serum (Gibco), and 1% antibiotic-antimycotic (Gemini Biosciences). MCF10A cells (ATCC) were cultured at 37°C with 5% CO2 in DMEM-F12 (Gibco) with 0.1% insulin (Sigma), 0.05% hydrocortisone (Sigma).

The following antibodies and stains were used for immunofluorescence imaging: laminin anti-mouse (Abcam) and AlexaFluor-555 goat anti-mouse (Life Technologies), AlexaFluor-647 phalloidin (Life Technologies) to label F-actin, and DAPI (Invitrogen) to label nuclei.

Laminin for cellular protein patterning was purchased from Sigma-Aldrich. Force disruption experiments utilized Calyculin A (Med Chem Express) and Rho-kinase inhibitor, Y27632 (Sigma-Aldrich).

### Image Processing and Analysis

Fluorescence image analysis was performed using custom MATLAB code, available upon request. Fluorescence images of both the hexago-where *i* spans the range of 6 neighboring spots. This process is iterated for all dots until a converged solution is reached in which 0.01% cholera toxin (Sigma), 1% antibiotic-antimycotic (Gemini Biosciences),nal laminin pattern and the actin cytoskeleton were imported into MATLAB, adjusted for intensity, and converted to binary images. Centroids for each spot in the hexagonal pattern were determined using the *regionprops* function in MATLAB (Fig. 3A). Binary images of the actin cytoskeleton were used to calculate cell size, and were superimposed onto the hexagonal pattern to identify cell locations within the pattern.

### Delaunay Triangulation and Identification of Undeflected Pattern Locations

A significant limitation of many cellular traction force measurement systems is the need to remove cells in order to determine the undeflected position of deflection markers. To overcome this limitation, we patterned surfaces with a hexagonal pattern of laminin circles. The hexagonal pattern was then used to generate a Delaunay triangulation, which created a triangle mesh from an array of points such that no point is contained within the circumcircle of any triangle (Fig. 3C). This Delaunay triangulation represents the deflected position of each triangular region of the substrate. To determine the undeflected position, we utilized the fact that the undeflected Delaunay triangles for a uniform hexagonal pattern should be equilateral triangles, and as such, each spot should be equidistant from its 6 neighboring spots (Fig. 3B). For each spot in the pattern, the centroid [*C*_*x*_ *C*_*y*_] is replaced with a new centroid given by:

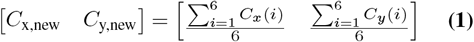

where i spans the range of 6 neighboring spots. This process is iterated for all dots until a converged solution is reached in which

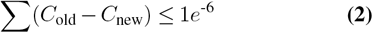

suggesting that all spots are now at a minimum distance from each of its six neighbors. The Delaunay triangulation of the new array of spots represents the undeflected positions (Fig. 3C), and the distance between the two triangulations gives the deflection vectors (Fig. 3D). One shortcoming of this method is error due to missing or misidentified spots, as commonly occurs in microcontact printing.

### Identifying and Replacing Missing or Misidentified Spots in the Hexagonal Pattern using Voronoi Diagrams

To avoid issues with missing or misidentified spots on the microcontact printed substrate, we generated Voronoi diagrams and looked for degenerate geometries. A Voronoi diagram is a dual graph of the Delaunay triangulation, in which a point is placed at the centroid of the circumcircle that defines each Delaunay triangle, and each point is connected to all neighboring points (Fig. 4A). For a hexagonal pattern, the Voronoi diagram should be a series of connected hexagons; if microcontact printed spots are absent in the pattern, the Voronoi spaces surrounding the missing dot will be degenerate, with either more or less than six sides. Additionally, the degenerate Voronoi regions should share a common vertex, which corresponds to the location of the missing printed spot. To replace missing or misidentified spots, we pooled all vertices that corresponded to degenerate Voronoi regions, and selected only vertices that were connected to two or more regions. These vertices were then added to the set of hexagonal pattern centroids.

### Calculating Plane Strain and Plane Stress in Triangular Elements

Once the deflected and undeflected positions of each triangle vertex were determined, these deflections were used to calculate the plane strain within the triangular element. Given that previous work has demonstrated that cells plated onto a 2D surface generate primarily in-plane forces (17), we used the finite element derivations for linear plane strain elements to calculate the 3 strains (x, y, and torsion) for each triangle:

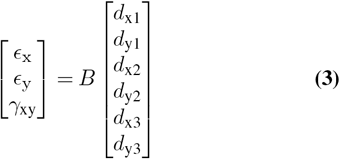

where

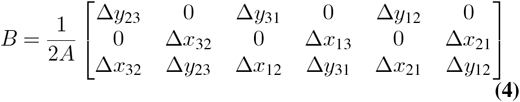

where A is the triangle area, [d] is the vector of deflections of the three triangle vertices, and Δx and Δy are the change in the x and y coordinates respectively of the triangle vertices indicated by subscript.

The 3-element strain vector is then used to calculate principal strains using the following transformations:

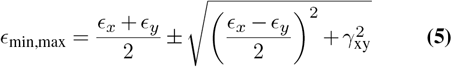

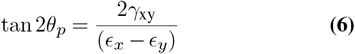

where max,min represent the maximum and minimum principal strains, and p is the angle that corresponds with the direction of principal strain.

This results in a determination of the maximum and minimum principal strains, as well as the angle that corresponds to the direction of principal strain. Fig. 4B shows a strain map of a representative cell, where each triangle is colored according to its maximum principal strain. Fig. 4C shows a vector map of principal strains from a representative cell, in which each principal strain is shown as a vector whose magnitude corresponds with maximum principal strain and whose direction corresponds with the angle of maximum strain. As we have previously observed in other traction force measurement systems, principal strains are directed towards the cell centroid (18).

Outlines of cells were determined using the actin image, which was overlaid on the deflection map as discussed above. In order to quantify strains surrounding the cell, the cell outline was dilated by approximately 10 *µ*m to include the relevant strained area surrounding the cells. It is worth noting that strains beneath the cell (Fig. 4B, black outline) are small, while the most substantial strains generated by the cell are in the area surrounding the cell (Fig. 4B, colored outlines). To calculate stresses within each triangle, we assumed that PDMS behaves as a purely elastic substrate. While PDMS is a viscoelastic material, the low strain rate of applied traction forces makes this assumption reasonable for the time scales at which we observed cellular traction forces. Plane stress in each triangle is calculated as:

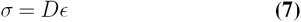

where

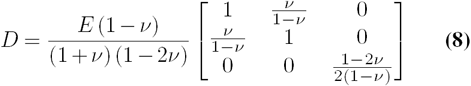

and E and *v* are the elastic modulus and the Poisson’s ratio of the substrate, respectively.

The total force in each triangle was then calculated by multiplying the maximum principal stress with the area of the Delaunay triangle. Total and average force per cell were calculated in two distinct ways, which are discussed below.

## Results

### Quantification of Average Strain and Average Stress Per Cell

As a proof of concept, MCF10A human mammary epithelial cells were treated with 2 ng/mL TGF-*β*1, which is known to increase traction forces in these cells (Fig. 5A-B). Cells were subsequently incubated with TGF-*β*1 in conjunction with 140 nM of the Rho-Kinase Inhibitor (Y27632), which blocks myosin contractility, or 2 nM Calyculin A, which has been shown to increase traction forces by blocking phosphorylation of Myosin Light Chain Phosphatase (MLCP). Total cell forces were determined by summing the forces under either: a) only the region surrounding the cell (Fig. 5C); or b) the entire area beneath and surrounding the cell (Fig. 5D). TGF-*β*1 significantly increased cellular traction forces surrounding each cell; inhibition of Rho Kinase blocked this increase, while inhibition of MLCP increased force (Fig. 5C). When forces underneath the cell were included in the analysis (Fig. 5D), similar trends were observed, with one exception: Calyculin A did not increase traction forces beyond the TGF-*β*1 condition. This can be attributed to the fact that most of the force generation occurs in the area surrounding the cell; inclusion of smaller forces underneath the cell thus reduces the difference between the TGF-*β*1 case versus the co-incubation of TGF-*β*1 and Calyculin A. In addition to altering cell forces, TGF-*β*1 also increases cell size. Interestingly, both Rho-Kinase inhibition and MLCP inhibition inhibited TGF-*β*1-induced cell spreading (Fig. 5E). As such, we sought to calculate the average cell stress, defined as the total cell-generated force per cell area (Fig. 5F). These data indicate that despite the increase in force, the cell stress decreases in epithelial cells that have been incubated with either TGF-*β*1 or Calyculin A.

**Fig. 5.**
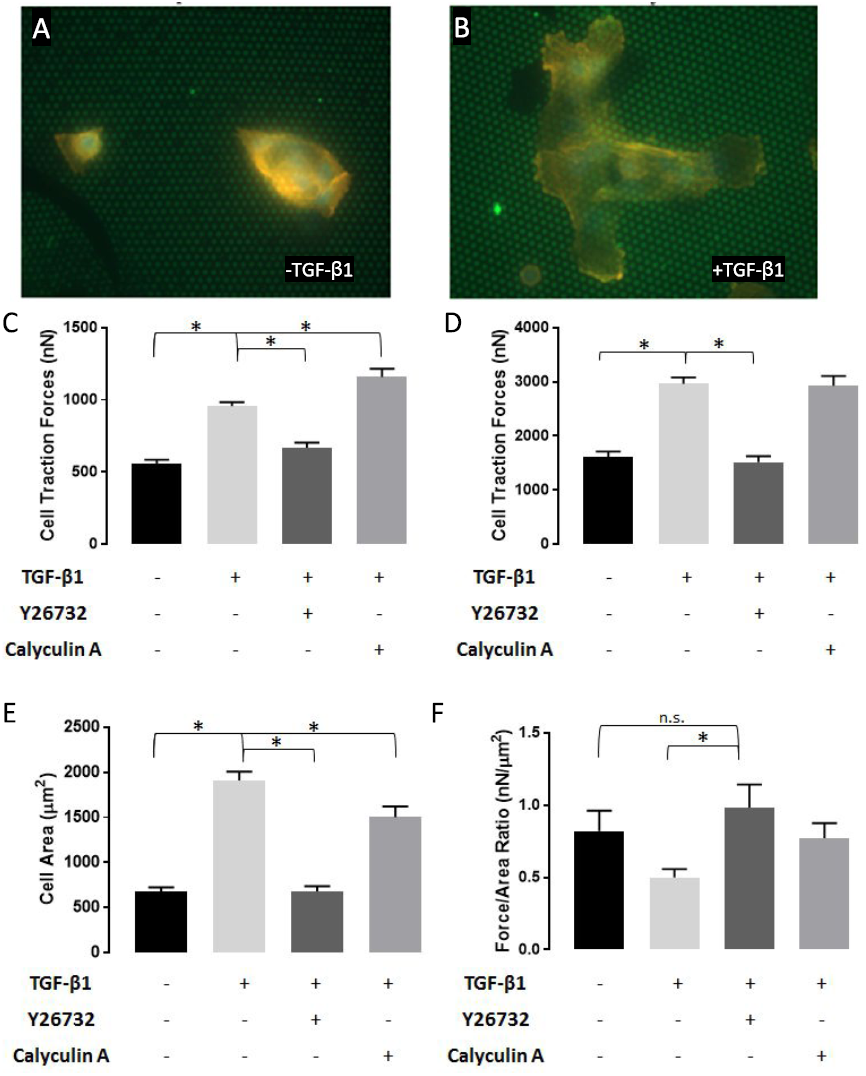
Comparison and Quantification of Traction Forces in MCF10A cells. A) Representative MCF10A cell on hexagonal pattern. B) Representative MCF10A cell treated with 2 ng/mL TGF-*β*1 on hexagonal pattern. C) Total cell traction force for area peripheral to cell outline. D) Total cell traction force for areas peripheral to and underneath cells. E) Average cell area F) Total cell force normalized to cell area. * indicates p-value of < 0.05, N > 15.

## Conclusions

Taken together, we have demonstrated a system that quantifies cell-generated traction forces in cells of low cytoskeletal rigidity, on a continuous substrate, without the need to remove cells. Data indicate that the system is capable of quantifying forces from epithelial cells, and the system captures force changes induced by TGF-*β*1. Positive and negative controls indicated that the system successfully detected changes in force when actomyosin force was increased or decreased, with the Rho-Kinase inhibitor Y27632 or Calyculin A, respectively.

## ACKNOWLEDGEMENTS

This work was supported by the National Science Foundation via Grant CMMI-1537168 (CAL, NAR) and by the National Institutes of Health/ National Institute of General Medical Sciences (CAL, BPG) via Grant R01-122855.

